# A peptide derived from the SARS-CoV-2 S2-protein heptad-repeat-2 inhibits pseudoviral fusion at micromolar concentrations: Role of palmitic acid conjugation

**DOI:** 10.1101/2023.05.13.540576

**Authors:** Nejat Düzgüneş, Zhihua Tao, Yuxia Zhang, Krzysztof Krajewski

## Abstract

SARS-CoV-2 S protein-mediated fusion is thought to involve the interaction of the membrane-distal, or N-terminal heptad repeat (NHR) (“HR1”) of the cleaved S2 segment of the protein, and the membrane-proximal, or C-terminal heptad repeat (CHR) (“HR2”) regions of the protein. Following the observations of Xia et al (Xia S, Liu M, Wang C, Xu W, Lan Q, Feng S, Qi F, Bao L, Du L, Liu S, Qin C, Sun F, Shi Z, Zhu Y, Jiang S, Lu L. Cell Res. 2020b Apr;30(4):343-355), we examined the fusion inhibitory activity of a PEGylated HR2-derived peptide and its palmitoylated derivative, using a pseudovirus infection assay. The latter peptide caused a 76% reduction in fusion activity at 10 μM. Our results suggest that small variations in peptide derivatization and differences in the membrane composition of pseudovirus preparations may affect the inhibitory potency of HR2-derived peptides.

## Introduction

The Severe Acute Respiratory Syndrome Coronavirus-2 (SARS-CoV-2) binds the angiotensin converting enzyme 2 (ACE2) receptor on its host cells via the S1 domain of its spike protein, S (Figure 1) (Düzgüneş et al., 2021). The transmembrane protease/serine subfamily (TMPRSS) cell surface proteases cleave the S protein into S1 and S2 sub-fragments. The receptor-binding domain (RBD) of S is located in the N-terminal S1 domain of S. Similar to the membrane fusion activity of the Env protein of HIV-1, the fusion activity of the S protein is associated with the C-terminal S2 domain.

**Figure 1.**
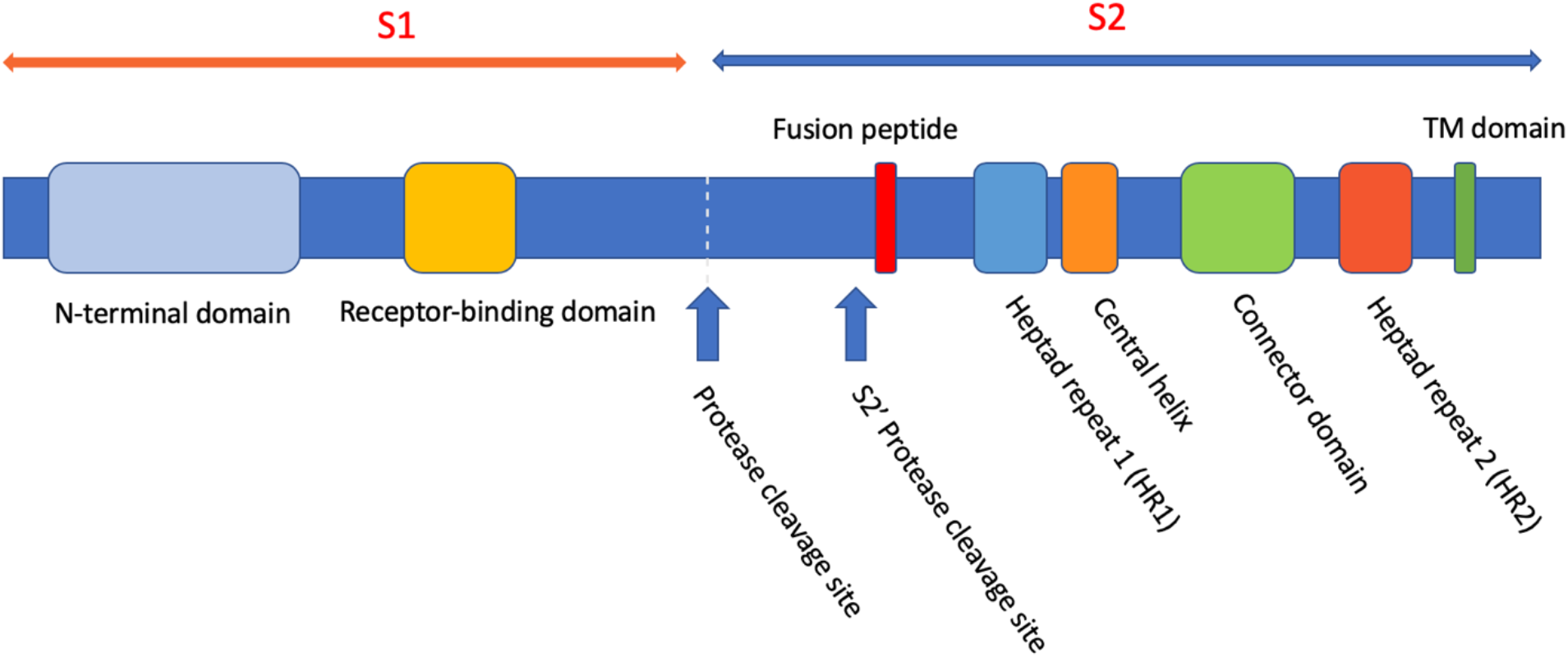
The domains of the SARS-CoV and SARS-CoV-2 spike protein, S. Reproduced from Düzgüneş & Konopka, 2020.

S-mediated fusion is thought to involve the interaction of the membrane-distal, or N-terminal heptad repeat (NHR) (also termed “HR1”) of S2, and the membrane-proximal, or C-terminal heptad repeat (CHR) (termed “HR2”) regions of the protein.

Upon binding to its receptor, the S-protein undergoes a conformational change where the S1 subunit is released, the fusion peptide of the S2 subunit is inserted into the host cell membrane, and the HR1 and HR2 domains interact with one another, resulting in the formation of the “six-helix bundle.” This conformational change brings the viral and cell membranes into close proximity (Figure 2) (Düzgüneş et al., 2021).

**Figure 2.**
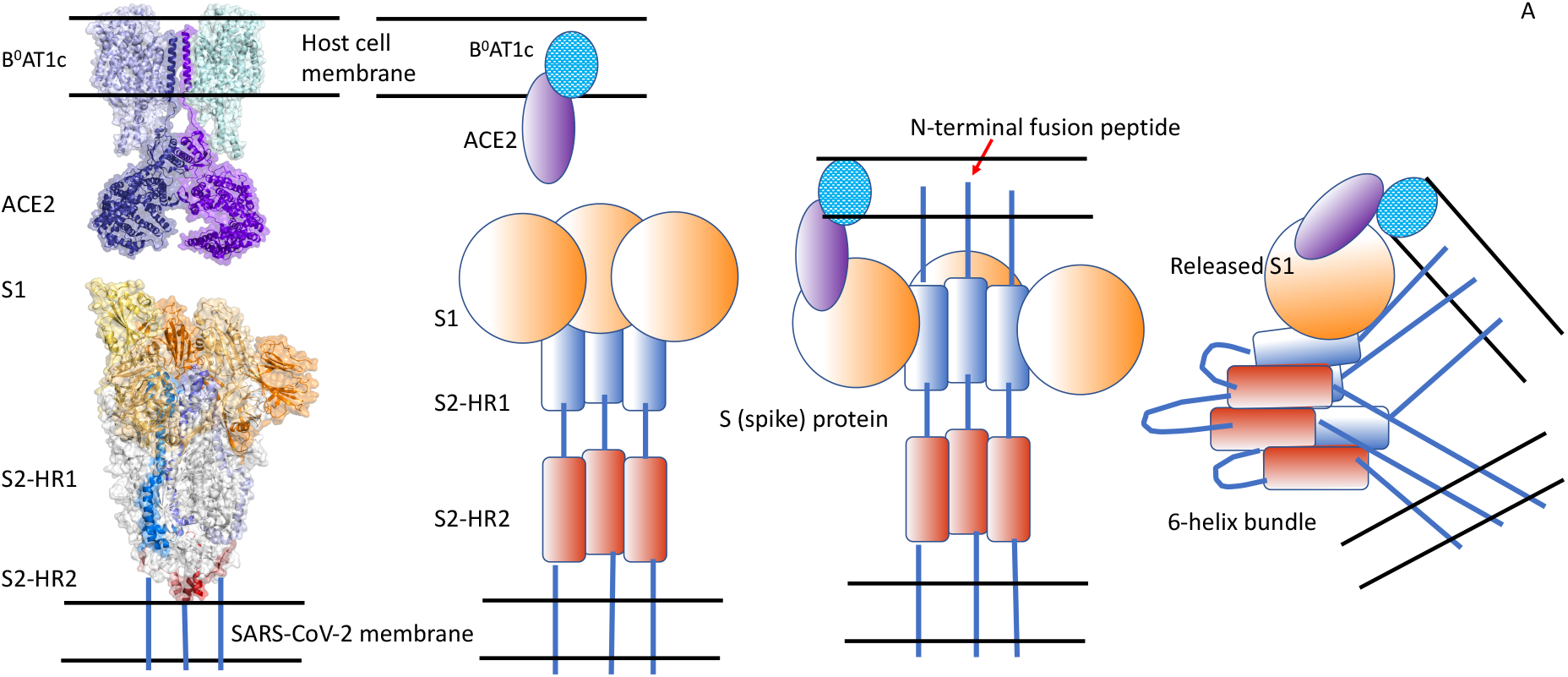
The conformational change of the SARS-CoV-2 spike protein and formation of the “six-helix bundle.” Reproduced from Düzgüneş et al., 2021.

The HR1 domain of SARS-CoV-2 has a high α-helicity and a high binding affinity to the HR2 domain. A peptide derived from the HR2 domain (“2019-nCoV-HR2P”) was found to inhibit cell-cell fusion induced by SARS-CoV-2 S-protein, with an IC_50_ of 0.18 μM (Xia et al., 2020a). These authors also showed that this peptide interacts with a peptide derived from the HR1 region and forms a helical structure characteristic of the 6-helix bundle, confirming the formation of the hairpin structure depicted in Figure 2. The same laboratory developed a similar peptide (“EK1”) derived from the HR2 region of the coronavirus OC43 and showed that it inhibits the fusion activity of multiple coronaviruses (Xia et al., 2019). They reported subsequently that EK1 inhibits SARS-CoV-2 cell-cell fusion at an IC_50_ of 0.32 μM (Xia et al. 2020b). They also found that linking cholesterol to the C-terminus of EK1 through 5 additional amino acids and poly(ethylene glycol)_4_ lowered the IC_50_ to 1.3 nM.

These observations indicate that synthetic peptides and their derivatives can be used as therapeutic agents against SARS-CoV-2. The sites of interaction of EK1 and possible other peptides with the S ptotein are shown in Figure 3.

**Figure 3.**
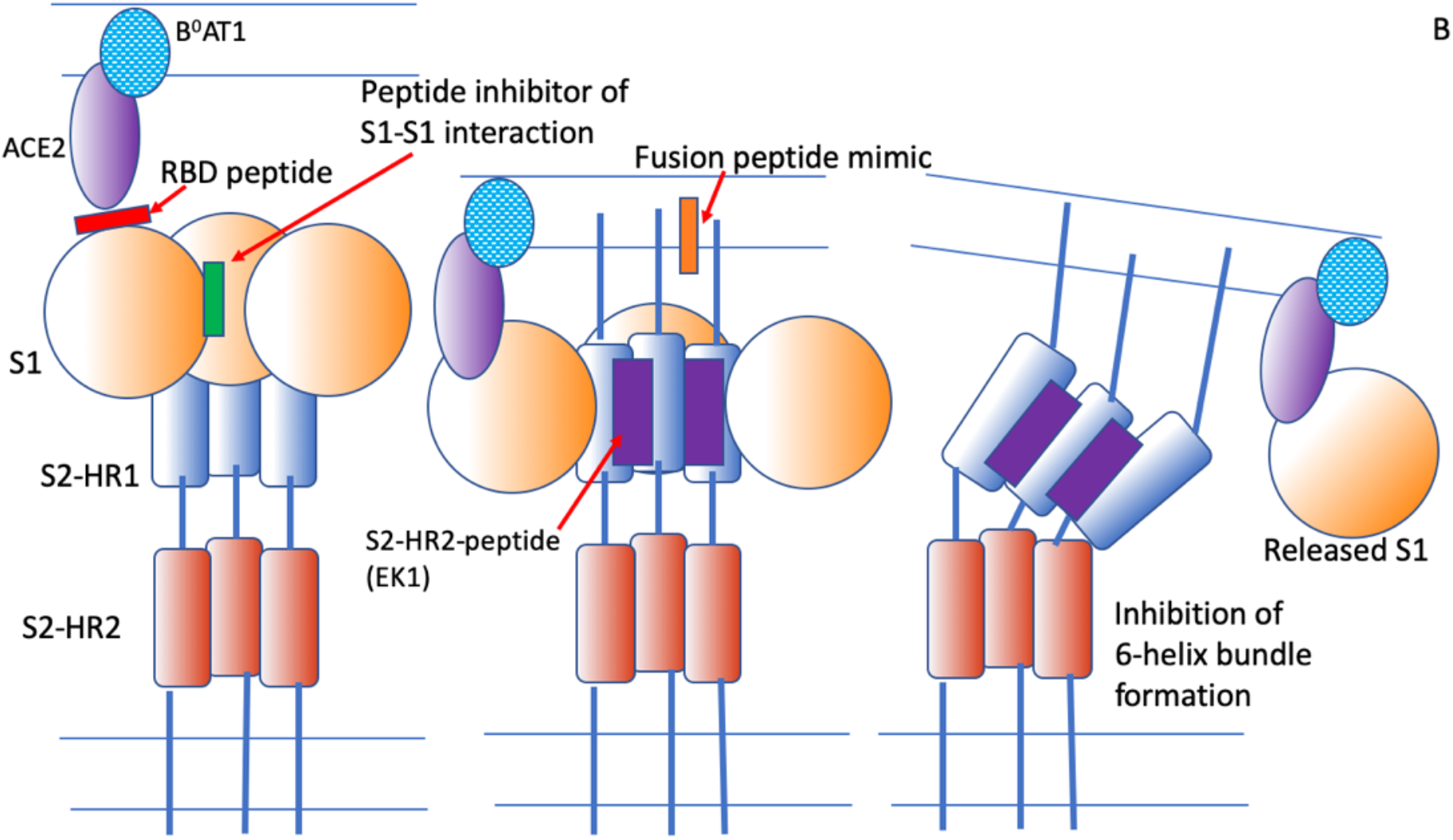
Inhibition of membrane fusion by various peptides. Red: A peptide that binds to the receptor-binding domain of S1. Green: A peptide that inhibits the interaction between S1 subunits. Orange: An S2 fusion peptide mimic that may inhibit the interaction of the fusion peptide with the target membrane. Purple: A peptide derived from the S2-HR2 region that binds with high affinity to the HR1 region and inhibits the interaction between the S2-HR1 and S2-HR2 domains, thus preventing “six-helix bundle” formation and membrane fusion. Reproduced from Düzgüneş et al., 2021.

Using a pseudotyped vesicular stomatitis virus expressing the SARS-CoV-2 S-protein, we tested the fusion inhibitory activity of a peptide identical to EK1, except for the addition of the PEG linker (PEG5) at the C-terminus (“Peptide 1”) and a palmitoyl derivative of Peptide 1, designated as “Peptide 2.”

## Materials and Methods

### Cells and supplies

The SARS-CoV2-Spike Pseudovirus and the ACE2-HEK293 Recombinant Cell Line were prepared in-house by BPS Bioscience; #79942 and, #79951, respectively) MEM medium (#SH30024.01), Na-pyruvate (#SH30239.01) and Pen-strep (#SV30010) were from Hyclone. Fetal bovine serum was obtained from Life

Technologies (#10082147). Non-essential amino acids (25-025-Cl) and 96-well tissue culture treated, white clear-bottom assay plates (#3610) were from Corning. Puromycin was purchased from InvivoGen (#ant-pr-1). The Spike S1 Neutralizing Antibody (Clone#G10xA1) (#101326) and the ONE-Step luciferase assay system (#60690) were from BPS Bioscience.

### Peptides

Peptides were synthesized at the UNC peptide synthesis facility (RRID:SCR_017837) on a CEM Liberty Blue microwave peptide synthesizer (HE-SPPS methodology, Collins et al., 2014) using Fmoc-ProTide Rink LL amide resin (loading 0.2 mmol/g). The C-terminal lysine residue was introduced with a ivDde sidechain protecting group; this allowed for orthogonal deprotection of the ε amino group of this residue (3 × 5min, 5% hydrazine in DMF), after the peptide sequence was fully synthesized and the N-terminal amine was protected with a Boc group. Subsequently the ε amino group of this Lys residue was coupled with Fmoc-NH-Peg_5_-COOH, and after Fmoc deprotection, for Peptide 2, coupled with palmitic acid. Both coupling reactions were performed manually using HATU as a coupling reagent in the presence of DIEA in DMF and confirmed with ninhydrin tests. The peptidyl resin was washed 3 x with dichloromethane, 3 x with methanol, and dried in a vacuum chamber overnight.

The peptide was cleaved from the resin and deprotected by a 2 h incubation with 2 mL of cleavage solution (2.5% trifluoroacetic acid (TFA), 2.5% TIPS, 2.5% EDT, 2.5% water) and precipitated by addition into cold diethyl ether (∼30 mL). The precipitate was collected by centrifugation and triturated twice with 5 mL of diethyl ether. Residual ether was allowed to evaporate, the peptide was dissolved in 50% acetonitrile and lyophilized. Crude peptides were purified by preparative RP HPLC on a Waters SymmetryShield RP18 column and lyophilized. Purified peptides were characterized by MALDI TOF MS and analytical HPLC (Millipore Chromolith RP-18e column).

### Cell culture and assay conditions

ACE2-HEK293 cells were cultured in MEM medium with 10% FBS, 1% non-essential amino acid, 1 mM Na-pyruvate, 1% Pen-strep, and 0.5 μg/mL puromycin. ACE2-HEK293 cells were seeded at 8,000 cells per well into white clear-bottom 96-well microplate in 90 μl of assay medium. Ten microliters of preincubated virus/compound mix were added into each well of the ACE2-HEK293 cells. For control cells, the same number of the ACE2-HEK293 cells were seeded, but no virus or compounds were added. The plates were incubated at 37°C and 5% CO_2_.

Approximately 48 h after transduction, the ONE-Step Luciferase reagent was prepared per the recommended protocol. One hundred microliters of One-Step Luciferase reagent were added per well and the plate was rocked at room temperature for ∼30 min. Luminescence was measured using a luminometer (BioTek Synergy™ 2 microplate reader).

### Data analysis

Reporter assays were performed in triplicate at each concentration. The luminescence intensity data were analyzed using the computer software, Graphpad Prism. In the absence of the compound, the luminescence intensity (L_t_) in each data set was defined as 100%. In the absence of pseudovirus, the luminescence intensity (L_b_) in each data set was defined as 0%. The percent luminescence in the presence of each compound was calculated according to the following equation: % Luminescence = (L–L_b_)/(L_t_– L_b_), where L= the luminescence intensity in the presence of the compound, L_b_ = the luminescence intensity in the absence of virus, and L_t_ = the luminescence intensity in the absence of the compound.

## Results and Discussion

The chemical composition of Peptide 1 is: SLDQINVTFLDLEYEMKKLEEAIKKLEES-YIDLKEL**K(Peg**_**5**_)-NH_2_ and that of Peptide 2 is:

SLDQINVTFLDLEYEMKKLEEAIKKLEES-YIDLKEL**K(Palm-Peg**_**5**_)-NH_2_.

Peptide 1 has the same amino acid sequence as the EK1 peptide of Xia et al. (2020b), except for the lysine linker at the C-terminus as well as a poly(ethylene glycol)_5_ (PEG5) moiety to render it a more appropriate control for the palmitoylated Peptide 2.

In our initial experiment we examined the effect of Peptide 1 and Peptide 2 on SARS-CoV-2 S-protein-mediated fusion. Peptide 1 and Peptide 2 had no inhibitory effect at concentrations of 0.1 to 1 μM (Figure 4). This result was surprising in view of the nanomolar inhibitory concentration of EK1 reported by Xia et al. (2020b) in an analogous system.

**Figure 4.**
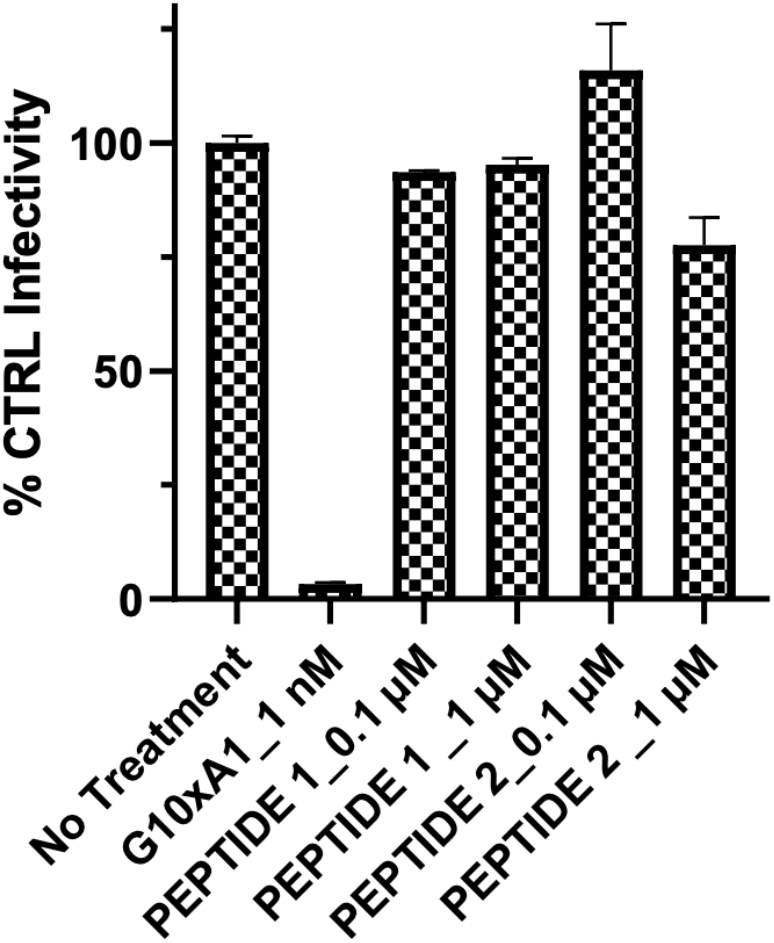
The effect of 0.1 μM and 1 μM Peptide 1 and Peptide 2 on the infectivity of the pseudotyped vesicular stomatitis virus expressing the SARS-CoV-2 S-protein.

We then examined higher concentrations of the peptides. Peptide 1 had 25% inhibitory activity at 10 μM, whereas Peptide 2 inhibited fusion by 76% at the same concentration (Figure 5).

**Figure 5.**
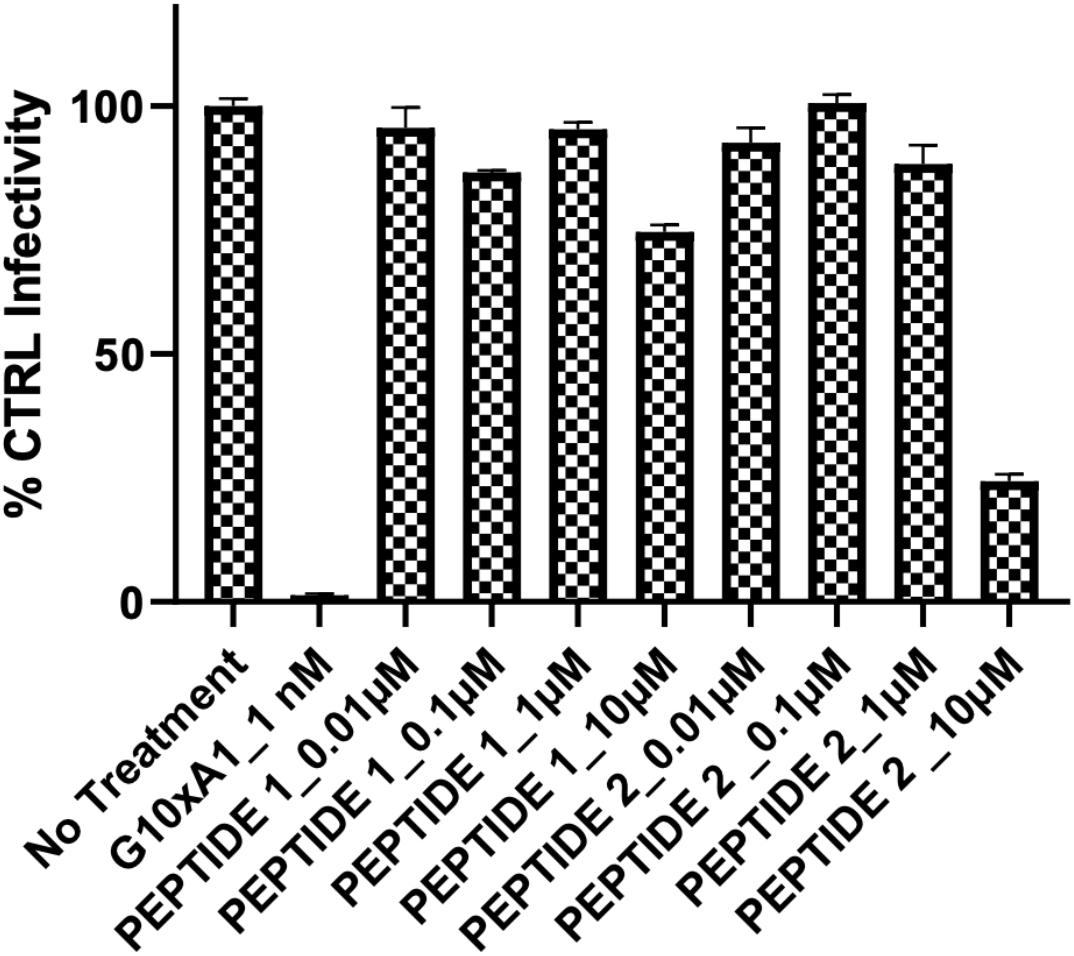
The effect of Peptide 1 and Peptide 2 in the range 0.01 μM to 10 μM on the infectivity of the pseudotyped vesicular stomatitis virus expressing the SARS-CoV-2 S-protein.

These inhibitory effects are much lower than those reported by Xia et al. (2020b). In an assay measuring SARS-CoV-2-mediated cell-cell fusion utilizing 293T/S/GFP effector cells and Huh-7 target cells, these authors found an IC_50_ of 287 nM for EK1 (corresponding to our Peptide 1) and an IC_50_ of 69 nM for EK1P (corresponding to Peptide 2). Their palmitoylated EK1P construct had a spacer of PEG4 instead of our PEG5. It is possible that small variations in the derivatization of the HR2-based peptides may affect the antiviral activity of the peptides.

In a pseudovirus fusion assay, the IC_50_ for EK1 was found to be 2.375 μM (Xia et al. 2020b); i.e. much higher than that found in the cell-cell fusion assay. Thus, it may be reasonable to expect that in our assay for SARS-CoV-2 S-pseudovirus fusion, inhibition of fusion by peptides may require higher concentrations than in a cell-cell fusion assay. Zhu et al, (2020) also observed much higher IC_50_s for their HR1-derived peptides in the pseudovirus assay than in their cell-cell fusion assay. It is also of interest to note that Xia et al (2020b) observed only a 60% inhibition of pseudovirus fusion by EK1 at 10 μM. Our Peptide 1, however, inhibited infectivity by only 25%.

In addition to the potential effects of small variations in peptide derivatization, it is likely that there are differences in the S-protein density in the pseudovirus membranes.

